# Etiology and antimicrobial resistance of bovine mastitis pathogens in North-Eastern Zimbabwe: Emerging threats and management gaps

**DOI:** 10.1101/2025.08.01.668064

**Authors:** Peter Katsande, Jianwen Liu, Min Zhou, Bamusi Saidi, Claudius Gufe, Crispen Rapangana, Addmore Waniwa, Tsamwi N Gurira, Shuvai Musari, Pious V Makaya, Chenai S Majuru

**Author notes:** Correspondence: Peter Katsande. These authors contributed equally to this work and share first authorship (Peter Katsande, Jianwen Liu).

## Abstract

Bovine mastitis, a significant challenge for dairy farming in Zimbabwe, affects milk production, animal health, and economic viability. This study investigated the etiology, prevalence and antimicrobial resistance of mastitis pathogens in dairy farms in the north-eastern regions of Zimbabwe. Of the 256 cows tested, 36.3% showed signs of mastitis, predominantly subclinical (28.1%). Key pathogens isolated included *Staphylococcus spp*. (34.4%), *Streptococcus spp*. (26.9%), and *Escherichia coli* (18.3%). Notably, *Candida spp*. and *Corynebacterium bovis* were identified for the first time as mastitis pathogens in Zimbabwe. The detection of *Candida spp*. underscores the impact of prolonged antibiotic use, which disrupts bacterial flora and promotes fungal infections, while the presence of *Corynebacterium bovis* highlights hygiene deficiencies in milking practices. Alarmingly, 25% of *Staphylococcus* isolates were multidrug-resistant (MDR), including methicillin-resistant *Staphylococcus aureus* (MRSA). Resistance to commonly used antimicrobials such as tetracycline and trimethoprim-sulfamethoxazole was widespread, further complicating treatment options. This study emphasizes the need for improved hygiene practices, routine screening, and antimicrobial stewardship to manage mastitis and curb the spread of resistance.

## Introduction

Bovine mastitis is the leading endemic infectious disease affecting dairy cows globally (Food and Agriculture Organization, 2014; Abebe et al, 2016). In Africa, particularly Zimbabwe, where there is limited research on mastitis, this disease is highly problematic. Mastitis causes significant economic losses to dairy farmers and the milk processing industry, due to decreased milk yield, discarded milk, changed milk composition, increased treatment costs, and veterinary expenses (Petrovski et al, 2006). Beyond its economic impact, mastitis poses a serious zoonotic risks and has been linked to the rising development and rapid emergence of multidrug-resistant strains worldwide (Pol and Ruegg, 2007; Oliver et al, 2011; Beyene et al, 2017).

Mastitis, an inflammatory response of the under tissue in the mammary gland, is typically caused by the adhesion, invasion and colonisation by mastitis pathogens occurs in three forms, namely clinical, sub-clinical, and chronic mastitis (Ruegg, 2017; Taponen et al, 2017). Clinical mastitis, which is less common, is marked by systemic symptoms in the cow as well as visible abnormalities in both the udder and milk (Jamali et al, 2018). In contrast, subclinical mastitis is more widespread and leads to reduced milk production without any noticeable clinical signs or abnormalities in the udder or milk (Ndahetuye et al, 2019). Due to the absence of clear signs, it is challenging to diagnose and tends to persist longer within the herd and is associated with greater economic losses compared to clinical mastitis (Abrahmsén et al 2013).

Milk processing and marketing organizations prefer milk with low somatic cell count (SCC) (<200,000cells/mL), and farms maintaining low SCC levels sell their milk at a premium price. However, many farmers struggle to reduce SCC levels on their farms. Cows with high SCC (>200,000 cell/mL) are most likely to have a high bacterial load, suffer from subclinical mastitis, and act as a reservoir of pathogenic bacteria, posing a risk to healthy cows. Farmers often fail to detect cows with high SCC, allowing them to remain in lactation without treatment.

The etiology of mastitis includes both contagious microorganisms, which survive and proliferate on the skin and teat wounds, and environmental microorganisms, which do not persist on the teat (Ruegg, 2017; Zeryehun et al, 2017). Mastitis is primarily caused by bacteria, and over 140 different pathogenic species have been identified globally (Motaung et al, 2017). Studies show significant variation in the prevalence and distribution of mastitis and its pathogens across different countries, regions, and farms, influenced by farm management practices and environmental factors (Verbeke et al, 2014; Gao, et al 2017 et al). Major mastitis pathogens have been already identified include Coliforms, *Streptococcus agalactiae*, and *Staphylococcus aureus* (Bradley, et al 2007; Zadoks, et al 2007).

Effective prevention and control of mastitis depend on timely detection and understanding of the diversity of pathogens associated with the disease (Bi, et al 2016). However, given the rapidly increasing antibiotic-resistant pathogens, treatment is expected to become increasingly challenging in the near future (Vakkamäki, et al 2017). Monitoring antimicrobial resistance (AMR) patterns in bacterial isolates from mastitis cases and farm environments is crucial not only for guiding treatment decisions but also for identifying potential reservoirs of resistant genes in dairy farms.

Several studies, have reported an increase in subclinical mastitis in the African region (Abrahmsén et al 2013; Zeryehun and Abera, 2017; Ndahetuye et al, 2019; Suleiman et al, 2018). However, research on the prevalence and etiology of mastitis in Zimbabwe remains limited. The only data that is available is from studies that were conducted more than a decade (Perry et al 1987; Makaya, Aarestrup and Olsen 1996; Kudinha and Simango 2002; Katsande et al 2013). This study, therefore, aimed to identify bacterial pathogens associated with clinical and subclinical mastitis on dairy farms in North-Eastern Zimbabwe and to determine their antimicrobial resistance profiles.

## Materials and Methods

### Study Farms and animal selection

The study was conducted from February to July 2024 on five dairy farms located in Bindura, Goromonzi and Marondera districts of north-eastern Zimbabwe. The study involved mixed breed cows, including Holstein, Holstein-Friesian, and Jersey. The cows varied in age, number of milking days, number of calves, and milk yield. Milking was conducted twice daily (morning and afternoon) using milking bucket-type machines. The cows were mainly fed fresh grass, wheat straw, and brans. Farms were selected based on monthly bulk tank somatic cell count (BTSCC) data from Dairy Services, with those exceeding 200,000 cells/mL being enrolled through voluntary participation of farmers. After selecting participating farms individual cows were screened for SCC, and those with SCC greater than 200,000 cells/mL) were identified for further analysis.

### Samples Collection

The sample size for the study was determined using Thrusfield’s formula, (2005), based on an estimated prevalence of 21.1%, from the previous report by Katsande et al. (2013) with a 95% confidence level and a precision of 5%. Accordingly, 256 lactating cows were included in the study, from which 180 milk samples were collected from cows showing clinical or subclinical mastitis.

Clinical mastitis was defined as an infection in one or more udder quarters exhibiting visible signs of inflammation such as redness, swelling, pain, increased heat, or visible changes in milk (e.g., watery, bloody, blood-tinged, serum-like) or consistency (e.g., clots or flakes or stringy/viscous texture). Subclinical mastitis was defined as an infection in one or more udder quarters of a cow without visible signs of inflammation in the udder or changes in milk but with an elevated SCC exceeding 200,000 cells/mL), determined by California Mastitis Test (CMT).

The SCC was determined at the Dairy Services Laboratory using the Soma Count 300 (Bentley Instruments Inc). The CMT was performed by mixing 2mL of milk from each quarter of the udder with an equal volume of 3% CMT reagent (CMT, AHDB Dairy, UK) in a four-well paddle, stirring for 10–15 seconds, and recording the results within 20 seconds. The presence of thickening or gel formation indicated elevated somatic cell counts. Results were interpreted using a scoring system from 0 to 4: 0 for no reaction, 1 for trace, 2 for weakly positive (presence of sediment), 3 for distinctly positive (sediment and a slight increase in consistency), and 4 for strongly positive (complete coagulation of the sample) as described by Sargeant et al. (2001).

Approximately 10 mL of milk samples were aseptically collected from each quarter of the udder into sterile plastic tubes. Prior to collection each cow’s teat was pre-dipped in a betadine solution, dried with an individual paper towel, and the teat opening was scrubbed with 70% alcohol. Individual milk samples were collected after discarding the first six streams of milk to eliminate any contaminating bacteria from the teat canal. Additional farm data were recorded for each sample, including cow identification number, the quarter of the udder sampled, and the clinical or subclinical status of the quarter.

The samples were transported in cooler boxes on ice, maintaining a temperature approximately 4° C to the Central Veterinary Laboratory for bacterial isolation and identification. Upon arrival, the samples were either cultured immediately or stored in a refrigerator at 4 -8° C for up to 24 hours before culturing.

### Bacterial Isolation and Identification

Milk samples that tested positive with CMT and with SCC > 200,000 cells/mL), were subjected to microbiological analysis, following procedures outlined by Quinn et al. (2011). A loopful (approximatively 0.01 mL) of milk was inoculated onto MacConkey agar (Oxoid, Basingstoke, UK) and blood agar plates enriched with 5% defibrinated sterile sheep blood. The inoculated plates were incubated aerobically at 37 °C for 24–48 h. After incubation, colony morphology was observed and recorded. Samples yielding more than one colony type were classified as mixed cultures. The distinct colonies were then subcultured to obtain pure colonies, which were subsequently used for final identification.

The pure bacterial isolates were identified through a series of phenotypic tests, including Gram staining, catalase, indole, and oxidase tests. *Staphylococcus* species were identified by testing for free coagulase presence and clumping factor using rapid slide agglutination tests (Bactident Coagulase, Merck, Darmstadt, Germany), catalase production and mannitol fermentation. Suspected pathogenic Staphylococcus strains were further tested on Staphytec Plus (Oxoid, Basingstoke, UK).

*Streptococcus* spp were identified using the catalase test, hemolytic activity, aesculin hydrolysis on Edward’s media (Oxoid, Basingstoke, UK) and its growth ability on MacConkey agar. Within group differentiation was done using the CAMP test. Gram-negative bacteria were identified based on lactose fermentation on MacConkey agar, along with tests for motility, indole production, citrate utilization, triple sugar iron reaction, urease activity, and oxidase production.

Final confirmation of bacterial species was done using the API biochemical profiling system, including API 20E, API Staph, API 20Strep, API NE, and API Coryne (bioMérieux, Marcy l’Etoile, France).

### Antimicrobial Susceptibility testing

Antimicrobial susceptibility test (AST) was performed using VITEK® 2 XL (bioMérieux version 08.01). AST for Gram-positive bacteria was performed using Vitek® 2 AST-GP67-22226 (bioMérieux, Marcy-l’Etoile, France) test card for Gram-positive susceptibility, AST-ST02 420915 Streptococcus and for Gram-negative bacteria, the Gram-negative susceptibility card (AST-GN81) were utilized. Bacterial suspensions were prepared in 0.45% sodium chloride solution, adjusted to a turbidity of 0.5 McFarland. The preparation and loading of the inoculum into the cards were completed within 30 minutes, after which the cards were placed in the Vitek-2 machine for incubation and automated reading.

The MIC interpretive standard or breakpoint values were set following the guidelines of Clinical and Laboratory Standard Institute M100-S23 (CLSI, 2024). Quality control was performed using *S. aureus* ATCC 25923 and *E. coli* ATCC 25922 as control strains. All susceptibility results for quality control strains were within the specified ranges.

## Results

### Microbiological Results

A total of 256 cows from 5 farms were tested for mastitis. Of these, 36.3% (93/256) had mastitis, with 28.1% (72/256) and 8.2% (21/256) having sub-clinical and clinical mastitis respectively. From the mastitis-positive cows, the most commonly isolated genera were *Staphylococcus* spp. (34.4%); *Streptococcus* spp (26.9%); *E. coli*, (18.3%); *Enterococcus faecium* (7.5%); *Corynebacterium bovis* (4.3%); *Enterobacter cloacae* (3.2%), *Candida* spp (3.2%), and *Klebsiella aerogenes* (2.2%) (Table 1). Among the *Staphylococcus* isolates, coagulase-positive staphylococci (CPS) accounted for 87.5% of all *Staphylococcus* species identified, with *S. aureus* comprising 71.9% and *S. intermedius* 15.6%. Coagulase-negative staphylococci included *S. epidermidis* (9.4%) and *S. xylosus* (3.1%). Within the Streptococcus group, *Streptococcus agalactiae* was the most prevalent species, accounting for 60% of isolates, followed by *Streptococcus dysgalactiae* (24%) and *Streptococcus uberis* (16%).

**Table 1.**
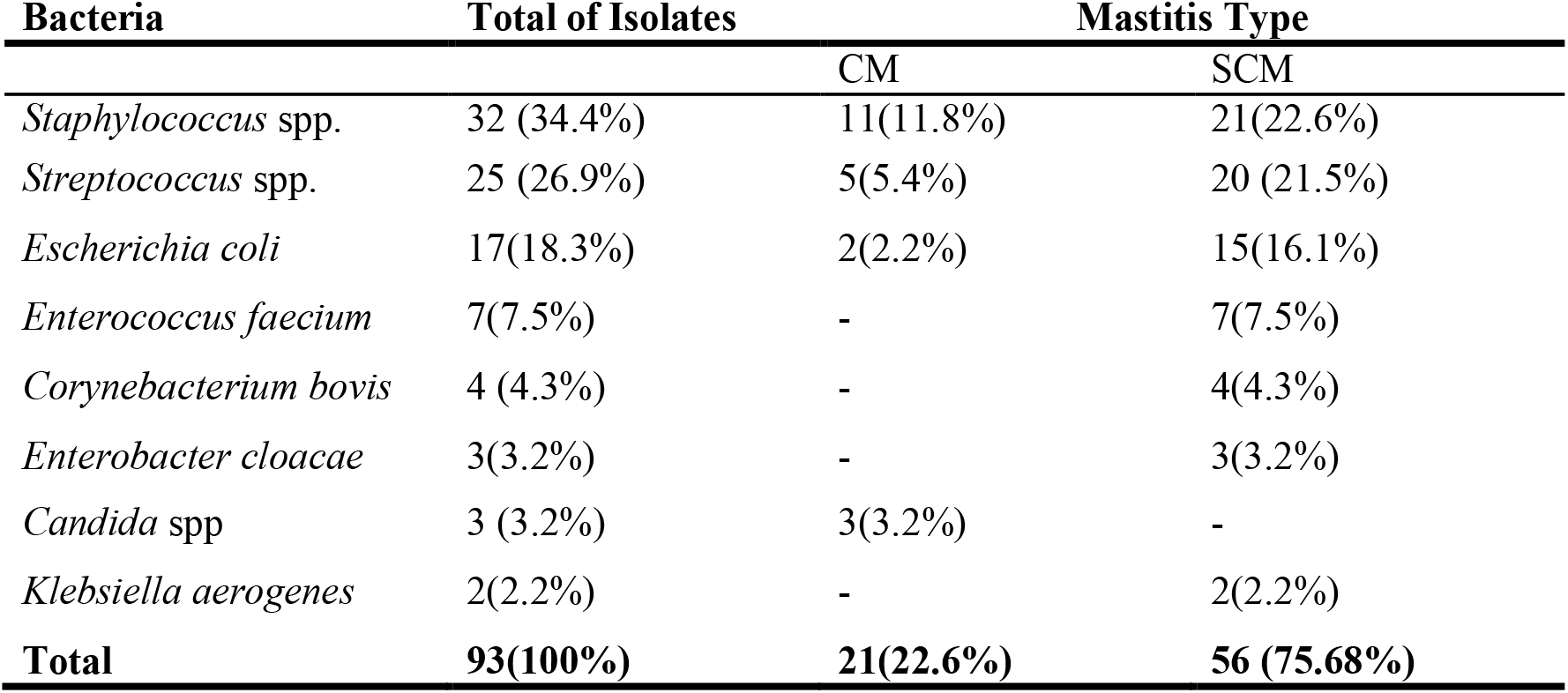
Microorganisms isolated from mastitic milk and their distribution depending mastitis type.

**Table 2.** Antimicrobial drug resistance profile of the isolated Staphylococci strains.

| Antimicrobial | *S. aureus*<br>(n=23) | *S. intermedius*<br>(n=5) | *S. epidermidis*<br>(n=3) | *S. xylosus*<br>(n=1) |
| --- | --- | --- | --- | --- |
| Benzylpenicillin | 8 | 1 | 0 | 0 |
| Oxacillin | 2 | 1 | 0 | 0 |
| Ciprofloxacin | 3 | 1 | 1 | 0 |
| Vancomycin | 2 | 1 | 0 | 0 |
| Erythromycin | 5 | 1 | 2 | 1 |
| Tetracycline | 12 | 3 | 0 | 0 |
| Trimethoprim/Sulfamethoxazole | 12 | 2 | 3 | 0 |
| Cefoxitin Screen | 2 | 1 | 0 | 0 |

### Antimicrobial Susceptibility Assay

All *Staphylococcus* spp. strains (n = 32) exhibited resistance to at least one antimicrobial agent. Notably, eight strains (25%) were classified as multidrug-resistant (MDR), showing resistance to three or more antimicrobial classes. The highest levels of resistance were observed for trimethoprim/sulfamethoxazole (53.1%, n=17) and tetracycline (46.9%, n=15). Low resistance was recorded for ciprofloxacin (15.6%, n=5), and vancomycin (9.4%, n=3). No resistance was detected for clindamycin, levofloxacin, linezolid, moxifloxacin, nitrofurantoin, quinupristin/dalfopristin and rifampicin. Additionally, three *S. aureus* isolates were identified as methicillin-resistant, based on their resistance to oxacillin and cefoxitin.

Streptococci strains were resistant to 5 out of the 13 antimicrobials tested (Table 3). Specifically, resistance was observed to tetracycline (40%, n=10), erythromycin (32%, n=8), trimethoprim/sulfamethoxazole (32%, n=8), penicillin (20%, n=5), and ampicillin (12%, n=3). Notably, two strains exhibited multidrug resistance, being resistant to three or more antibiotics. However, all strain were susceptible to cefotaxime, ceftriaxone, clindamycin, levofloxacin, linezolid, moxifloxacin. For Enterococcus strains, resistance was observed only to ampicillin (14.3%, n=1) and penicillin (14.3%, n=1).

**Table 3.** Antimicrobial drug resistance profile of the isolated streptococci strains.

| Antimicrobial | *S. agalactiae*<br>(n=17) | *S. dysgalactiae*<br>(n=5) | *S. Uberis*<br>(n=3) | *E. faecium*<br>(n=7) |
| --- | --- | --- | --- | --- |
| Ampicillin | 3 | 0 | 0 | 1 |
| Benzylpenicillin | 2 | 1 | 2 | 1 |
| Erythromycin | 5 | 2 | 1 | 0 |
| Tetracycline | 3 | 5 | 2 | 0 |
| Trimethoprim/Sulfamethoxazole | 5 | 3 | 0 | 0 |

**Table 4.** Antimicrobial drug resistance profile of the isolated *E*.*coli, K. aerogenes* and *E. cloacae* strains.

| Antimicrobial | *E.coli*<br>(n=17) | *K. aerogenes*<br>(n=2) | *E. cloacae*<br>(n=3) |
| --- | --- | --- | --- |
| Ampicillin | 9 | 1 | 1 |
| Cefazolin | 2 | 0 | 0 |
| Cefoxitin | 2 | 0 | 0 |
| Cefepime | 2 | 0 | 0 |
| Ceftazidime | 2 | 0 | 0 |
| Ceftriaxone | 2 | 0 | 0 |
| Ciprofloxacin | 2 | 0 | 0 |
| Nitrofurantoin | 2 | 0 | 0 |
| Tetracycline | 4 | 1 | 2 |
| Trimethoprim/Sulfamethoxazole | 9 | 1 | 3 |
| ESBL | 2 | 0 | 0 |

*E. coli* strains demonstrated resistance to ampicillin and trimethoprim/sulfamethoxazole (52.9%, n=9), tetracycline (23.5%, n=4), nitrofurantoin (11.8%, n=2), and to cefazolin, cefoxitin, cefepime, ceftazidime, ceftriaxone, and ciprofloxacin (11.8%, n=2 each). Two of the *E*.*coli* isolates were ESBL. However, all *E. coli* strains were susceptible to amoxicillin/clavulanic acid, piperacillin/tazobactam, meropenem, amikacin, and gentamicin.

*K. aerogenes* was only resistant to 3 antibiotics, ampicillin, tetracycline and trimethoprim/sulfamethoxazole. *E. cloacae* exhibited resistance to ampicillin, tetracycline, and trimethoprim/sulfamethoxazole.

## Discussions

The overall mastitis prevalence of 36.3% aligns with previous studies in sub-Saharan Africa, where rates range between 20–40% depending on management practices and diagnostic methods (Katsande et al; Abera et al., 2013). Sub-clinical mastitis (26.2%) was far more prevalent than clinical mastitis (2.7%), consistent with earlier studies highlighting sub-clinical mastitis as a hidden yet significant cause of economic losses in dairy production (Tolosa and Tigre 2009; Mbindyo et al., 2021). This is likely partly influenced by the chronic nature of subclinical mastitis, where cases persist longer than clinical ones and accumulate over time, as well as by the economic focus of producers, who often prioritise treating clinical cases due to their visible impact on milk yield and cow health, leaving subclinical cases unnoticed and untreated.

The high prevalence of sub-clinical mastitis observed in this study could possibly indicate considerable economic losses, primarily through reduced milk production caused by the progressive destruction of alveolar epithelial cells in the mammary gland (Zhao and Lacasse, 2008). Similar findings were reported by Mungube et al. (2005), who highlighted the financial burden of sub-clinical mastitis. This high prevalence underscores the need for regular screening and improved management practices in Zimbabwean dairy farms.

The predominant bacterial genera isolated were *Staphylococcus spp*. (34.4%), *Streptococcus spp*. (26.9%), and *Escherichia coli* (18.3%), corroborating global reports where these pathogens are the leading etiological agents of mastitis (Bradley, 2002; Poutrel et al, 2018; Pascu et al, 2022). Coagulase-positive staphylococci (CPS), particularly *S. aureus*, were the most prevalent among staphylococcal isolates. The dominance of *S. aureus* is consistent with its role as a major pathogen in bovine mastitis due to its ability to invade mammary tissue and evade host defences (Fagundes et al., 2010). The detection of *S. epidermidis* and *S. xylosus* highlights the potential role of coagulase-negative staphylococci (CNS) as emerging mastitis pathogens. *Streptococcus agalactiae* was the most commonly identified species among the streptococci, consistent with its well-documented pathogenicity in bovine mastitis (Hernandez et al, 2021; Tomazi et al, 2018; Lakew et al, 2019). *E. coli* accounted for 16.2% of isolates and is a well-known environmental mastitis pathogen. The presence of *Enterobacter cloacae* (5.41%) and *Klebsiella aerogenes* further emphasizes the importance of environmental hygiene in mastitis prevention.

The detection of *Candida spp*. and *Corynebacterium bovis* as causative agents of mastitis in Zimbabwe highlights emerging challenges in dairy herd management. Fungal infections, such as those caused by *Candida spp*., are rare in bovine mastitis but are often associated with immunosuppression, prolonged antibiotic use, or unhygienic milking practices that introduce yeast into the mammary gland. These infections frequently follow inappropriate or extended antibiotic use, which disrupts the natural bacterial flora, allowing opportunistic fungi to thrive (Motaung et al., 2017). This points to potential antibiotic misuse or resistance in Zimbabwean dairy farms.

The identification of *Corynebacterium bovis*, commonly found on the skin or teat canal, as a mastitis pathogen suggests a shift in pathogen profiles, with less aggressive organisms exploiting weakened udder defences. This indicates suboptimal hygiene practices during milking (Karzis, 2020). The first-time recognition of these pathogens in Zimbabwe emphasises the need for national pathogen surveillance to better understand local mastitis dynamics. It also underscores the importance of updating treatment protocols to address non-bacterial pathogens such as *Candida spp*.

All *Staphylococcus* isolates exhibited resistance to at least one antimicrobial agent, with 25% being multidrug-resistant (MDR). The high resistance to trimethoprim/sulfamethoxazole (and tetracycline aligns with prior studies that report these agents as overused in veterinary practice (Kadlec et al., 2015). Alarmingly, three *S. aureus* isolates were identified as methicillin-resistant (MRSA), indicating a potential zoonotic threat (Watts et al., 2018).

However, the susceptibility of all isolates to clindamycin, linezolid, and rifampicin provides alternative treatment options. Resistance to tetracycline and erythromycin was notable among streptococci. The emergence of MDR strains within this group warrants attention, as resistance to penicillin challenges the traditional use of this antibiotic for streptococcal infections.

Among *E. coli* isolates, resistance to ampicillin and trimethoprim/sulfamethoxazole was the highest. This finding mirrors reports from similar studies in sub-Saharan Africa (Karimuribo et al., 2006). The detection of extended-spectrum beta-lactamase (ESBL)-producing *E. coli* isolates further highlights the potential for resistance gene transfer in the dairy farm environment. However, all isolates were susceptible to critical antimicrobials like meropenem, piperacillin/tazobactam, and amikacin, providing effective therapeutic options.

The findings highlight critical gaps in antimicrobial stewardship and mastitis management in Zimbabwe. The high prevalence of MDR pathogens, including MRSA and ESBL-producing *E. coli*, poses significant threats to both animal and public health. Previous studies have identified a link between the misuse of antimicrobials in livestock and the emergence of resistant pathogens (Van Boeckel et al., 2017).

Improved farm hygiene, routine screening, and judicious use of antimicrobials are essential to mitigate the spread of resistant pathogens. Furthermore, surveillance programs should be established to monitor resistance trends, ensuring the sustainability of available treatment options.

## Conclusion

This study demonstrates a significant burden of bovine mastitis in Zimbabwe, with notable resistance patterns in the pathogens isolated. The emergence of MDR strains and zoonotic pathogens like MRSA underscores the urgent need for integrated approaches to mastitis control, incorporating better farm management, targeted therapy, and antimicrobial stewardship programs. Future research should explore the genetic mechanisms underlying resistance in these pathogens to inform evidence-based interventions.

## Supporting information

Supplementary Metadata and AST results

## Acknowledgements

The authors are grateful to the Department of Veterinary Technical Services and the International Exchange Service Center of the Ministry of Agriculture and Rural Affairs, Beijing, China for supporting the development of the original ideas for this project.

## Author contributions

Conceptualization: P.K. and J.L, Methodology and investigation: P.K., J.L., E.W., C. R., J.M., C.G., T.N.G., P.V.M., C.S.M. and B.S.

Data curation and formal analysis: P.K., J.L. and C.G. Writing – original draft preparation: P.K. and J.L

All authors read and approved the final manuscript. P.K. and J.L contributed equally to the manuscript and share first authorship.

## Ethical statement

Milk samples for this study were collected as part of routine veterinary diagnostic procedures for mastitis in dairy cattle, with the informed consent of the farm owners. According to current regulations and customary practices in Zimbabwe, the collection of milk from animals for mastitis screening does not require formal ethical approval from an institutional animal ethics committee, as it falls within the scope of standard veterinary care. No invasive procedures or experimental interventions were performed on the animals.

## References

Abebe R., Hatiya H., Abera M., Megersa B., and Asmare K., Bovine mastitis : prevalence, risk factors and isolation of Staphylococcus aureus in dairy herds at Hawassa milk shed, South Ethiopia, BMC Veterinary Research. (2016) 12, no. 1, 10.1186/s12917-016-0905-3, 2-s2.0-85000979429.

Abera, M., Habte, T., Aragaw, K., et al. (2013). Bovine mastitis: Prevalence, risk factors and major pathogens in the Ethiopian highlands. Tropical Animal Health and Production, 45(7), 1505–1511.

Abrahmsén, M.; Persson, Y.; Kanyima, B.M.; Båge, R. Prevalence of subclinical mastitis in dairy farms in urban and peri-urban areas of Kampala, Uganda. Trop. Anim. Health Prod. 2013, 46, 99–105

Beyene T., Hayishe H., Gizaw F. et al., Prevalence and antimicrobial resistance profile of Staphylococcus in dairy farms, abattoir, and humans in Addis Ababa, Ethiopia, BMC Research Notes. (2017) 10, no. 1, 1–9, 10.1186/s13104-017-2487-y, 2-s2.0-85018185639.

Bi Y., Wang Y. J., Qin Y., Vallverdú R. G., and García J. M., Prevalence of bovine mastitis pathogens in bulk tank milk in China, PLoS One. (2016) 11, no. 5, 1–13, e0155621, 10.1371/journal.pone.0155621, 2-s2.0-84969543226

Bradley A. J., Leach K. A., Breen J. E., Green L. E., and Green M. J., Survey of the incidence and aetiology of mastitis on dairy farms in England and Wales, Veterinary Record. (2007) 160, no. 8, 253–258, 10.1136/vr.160.8.253, 2-s2.0-33847738974.

Bradley, A. J. (2002). Bovine mastitis: An evolving disease. The Veterinary Journal, 164(2), 116–128.

Fagundes, H., & Oliveira, C. A. F. (2010). Pathogen-specific therapies in the control of bovine mastitis. Veterinary Microbiology, 144(1-2), 149–154.

Food and Agriculture Organization, Impact of mastitis in small scale dairy production systems, Animal Production and Health Working Paper, No. 13. (2014) Rome, Italy.

Gao J., Barkema H. W., Zhang L. et al., Incidence of clinical mastitis and distribution of pathogens on large Chinese dairy farms, Journal of Dairy Science. (2017) 100, no. 6, 4797– 4806, 10.3168/jds.2016-12334, 2-s2.0-85018626934.

Hernandez Luciana, Bottini Enriqueta, Cadona Jimena, Cacciato Claudio, Monteavaro Cristina, Bustamante Ana, Sanso Andrea Mariel (2021). Multidrug Resistance and Molecular Characterization of Streptococcus agalactiae Isolates From Dairy Cattle With Mastitis. Frontiers in Cellular and Infection Microbiology 11. URL= https://www.frontiersin.org/journals/cellular-and-infection-microbiology/articles/10.3389/fcimb.2021.647324. xDOI=10.3389/fcimb.2021.647324.

Jamali H., Barkema H. W., Jacques M. et al., Invited review: incidence, risk factors, and effects of clinical mastitis recurrence in dairy cows, Journal of Dairy Science. (2018) 101, no. 6, 4729–4746, 10.3168/jds.2017-13730, 2-s2.0-85042852702.

Kadlec, K., & Schwarz, S. (2015). Antimicrobial resistance of Staphylococcus aureus and Staphylococcus pseudintermedius in veterinary medicine. Microbiology Spectrum, 3(1).

Karimuribo, E. D., Fitzpatrick, J. L., Bell, C., et al. (2006). Clinical and subclinical mastitis in smallholder dairy farms in Tanzania: Risk, intervention and knowledge transfer. Preventive Veterinary Medicine, 74(2-3), 84–98.

Karzis, J. (2020). “Antibiotic Resistance in Dairy Herds. University of Pretoria (South Africa) ProQuest Dissertations & Theses, 2020. 31603922.

Katsande, S., Matope, G., Ndengu, M. and Pfukenyi, D.M., 2013, ‘Prevalence of mastitis in dairy cows from smallholder farms in Zimbabwe’, Onderstepoort Journal of Veterinary Research 80(1), Art. #523, 7 pages. 10.4102/ojvr.v80i1.523 (http://dx.doi.org/10.4102/ojvr.v80i1.523)

Katsande, Simbarashe et al. Prevalence of mastitis in dairy cows from smallholder farms in Zimbabwe. Onderstepoort j. vet. res., Pretoria, v. 80, n. 1, p. 00, Jan. 2013. Available from <http://www.scielo.org.za/scielo.php?script=sci_arttext&pid=S0030-24652013000100007&lng=en&nrm=iso>. access on 10 Jan. 2024

Kudinha, T. & Simango, C. 2002, ‘Prevalence of coagulase-negative staphylococci in bovine mastitis in Zimbabwe’, Journal of the South African Veterinary Association 73(2), 62–65. 10.4102/jsava.v73i2.557, PMid:12240771

Kudinha, T. and Simango, C. 2002, ‘Prevalence of coagulase negative staphylococci in bovine mastitis in Zimbabwe’, Journ of the South African Veterinary Association 73(2), 62– 65. 10.4102/jsava.v73i2.557 (http://dx.doi.org/10.4102/jsava.v73i2.557), PMid:122407

Lakew, B., Fayera, T. & Ali, Y. Risk factors for bovine mastitis with the isolation and identification of Streptococcus agalactiae from farms in and around Haramaya district, eastern Ethiopia. Trop Anim Health Prod 51, 1507–1513 (2019). 10.1007/s11250-019-01838

Makaya, P.V., Aarestrup, F.M. & Olsen, J.E., 1996, ‘Distribution and antibiotic resistance patterns of common mastitis pathogens (Gram-positive cocci) in selected dairy herds of three farming dairy sectors in Zimbabwe’, Zimbabwe Veterinary Journal 27(2), 65–75

Motaung T. E., Petrovski K. R., Petzer I.-M., Thekisoe O., and Tsilo T. J., Importance of bovine mastitis in Africa, Animal Health Research Reviews. (2017) 18, no. 1, 58–69, 10.1017/s1466252317000032, 2-s2.0-85020706254.

Motaung TE, Petrovski KR, Petzer I-M, Thekisoe O, Tsilo TJ. Importance of bovine mastitis in Africa. Animal Health Research Reviews. 2017;18(1):58–69. doi:10.1017/S1466252317000032.

Mungube, E.O., Tenhagen, B.A., Regassa, F., Kyule, M.N., Shiferaw, Y., Kassa, T. et al., 2005, ‘Reduced milk production udder quarters with subclinical mastitis and associated econo loss in crossbred dairy cows in Ethiopia’, Tropical Animal Heal and Production 37, 503–512. 10.1007/s11250-005-7049-y (http://dx.doi.org/10.1007/s11250-005-7049-y), PMid:16248222

Ndahetuye J. B., Persson Y., Nyman A.-K., Tukei M., Ongol M. P., and Båge R., Aetiology and prevalence of subclinical mastitis in dairy herds in peri-urban areas of Kigali in Rwanda, Tropical Animal Health and Production. (2019) 51, no. 7, 2037–2044, 10.1007/s11250-019-01905-2, 2-s2.0-85065036374.

Oliver S. P., Murinda S. E., and Jayarao B. M., Impact of antibiotic use in adult dairy cows on antimicrobial resistance of veterinary and human pathogens: a comprehensive review, Foodborne Pathogens and Disease. (2011) 8, no. 3, 337– 355, 10.1089/fpd.2010.0730, 2-s2.0-79952376274

Pascu, C.,, Herman, V.,, Iancu, I.,, Costinar, L. Etiology of Mastitis and Antimicrobial Resistance in Dairy Cattle Farms in the Western Part of Romania. Antibiotics 2022, 11, 57. 10.3390/antibiotics11010057.

Perry, B.D., Carter, M.E., Hill, F.W.G. & Milne, J.A.C., 1987, ‘Mastitis and milk production in cattle in a communal land of Zimbabwe’, British Veterinary Journal 143, 44–50. 10.1016/0007-1935(87)90105-

Petrovski, K.,, Trajcev, M.,, Buneski, G. A review of the factors affecting the costs of bovine mastitis: Review article. J. S. Afr. Vet. Assoc. 2006, 77, 52–60

Pol M. and Ruegg P. L., Relationship between antimicrobial drug usage and antimicrobial susceptibility of gram-positive mastitis pathogens, Journal of Dairy Science. (2007) 90, no. 1, 262–273, 10.3168/jds.s0022-0302(07)72627-9, 2-s2.0-35748950432

Poutrel B, Bareille S, Lequeux G, Leboeuf F (2018) Prevalence of Mastitis Pathogens in France: Antimicrobial Susceptibility of Staphylococcus aureus, Streptococcus uberis and Escherichia coli. J Vet Sci Technol 9: 522. doi: 10.4172/2157-7579.1000522.

Quinn, J.,, Markey, B.K.,, Leonard, F.C.,, FitzPatrick, E.S.,, Fanning, S.,, Hartigan, P.J. Veterinary Microbiology and Microbial Disease, 2nd ed.,, Wiley-Blackwell, A John Wiley & Sons Ltd. Publications: Ames, IA, USA, 2011.

Ruegg, P.L. A 100-Year Review: Mastitis detection, management, and prevention. J. Dairy Sci. 2017, 100, 10381–10397. [CrossRef]

Sargeant, J.,, Leslie, K.,, Shirley, J.,, Pulkrabek, B.,, Lim, G. Sensitivity and Specificity of Somatic Cell Count and California Mastitis Test for Identifying Intramammary Infection in Early Lactation. J. Dairy Sci. 2001, 84, 2018–2024

Sergeant, ESG, 2018. Epitools Epidemiological Calculators. Ausvet. Available at: http://epitools.ausvet.com.au.

Suleiman T. S., Karimuribo E. D., and Mdegela R. H., Prevalence of bovine subclinical mastitis and antibiotic susceptibility patterns of major mastitis pathogens isolated in Unguja island of Zanzibar, Tanzania, Tropical Animal Health and Production. (2018) 50, no. 2, 259– 266, 10.1007/s11250-017-1424-3, 2-s2.0-85030536146.

Taponen, S.,, Liski, E.,, Heikkilä, A.-M.,, Pyörälä, S. Factors associated with intramammary infection in dairy cows caused by coagulase-negative staphylococci, Staphylococcus aureus, Streptococcus uberis, Streptococcus dysgalactiae, Corynebacterium bovis, or Escherichia coli. J. Dairy Sci. 2017, 100, 493–503

Thrusfield M. Veterinary epidemiology. 2nd ed. London: Black well science Ltd.,, 2005. p. 182–98.

Tomazi T, de Souza Filho AF, Heinemann MB, Santos MVd (2018) Molecular characterization and antimicrobial susceptibility pattern of Streptococcus agalactiae isolated from clinical mastitis in dairy cattle. PLoS ONE 13(6): e0199561. 10.1371/journal.pone.0199561.

Vakkamäki, J.,, Taponen, S.,, Heikkilä, A.M.,, Pyörälä, S. Bacteriological etiology and treatment of mastitis in Finnish dairy herds. Acta Vet. Scand. 2017, 59, 33

Van Boeckel, T. P., Brower, C., Gilbert, M., et al. (2017). Global trends in antimicrobial use in food animals. Proceedings of the National Academy of Sciences, 114(3), 546–551.

Verbeke J., Piepers S., De Vliegher K., and Vliegher S. D., Pathogen-specific incidence rate of clinical mastitis in Flemish dairy herds, severity, and association with herd hygiene, Journal of Dairy Science. (2014) 97, no. 11, 6926–6934, 10.3168/jds.2014-8173, 2-s2.0-84908338784

Watts, J. L., & Salmon, S. A. (2018). Methicillin-resistant Staphylococcus aureus in dairy cattle: Epidemiology and control. Journal of Dairy Science, 101(5), 3938–3950.

Zadoks R. N. and Fitzpatrick J. L., Changing trends in mastitis, Irish Veterinary Journal. (2009) 62, no. 4, 59–70, 10.1186/2046-0481-62-s4-s59, 2-s2.0-70349949572

Zeryehun T. and Abera G., Prevalence and bacterial isolates of mastitis in dairy farms in selected districts of Eastern Harrarghe zone, Eastern Ethiopia, Journal of Veterinary Medicine. (2017) 2017, 7, 6498618, 10.1155/2017/6498618

Zeryehun, T.,, Abera, G. Prevalence and Bacterial Isolates of Mastitis in Dairy Farms in Selected Districts of Eastern Harrarghe Zone, Eastern Ethiopia. J. Vet. Med. 2017, 2017, 6498618.

Zhao, X. and Lacasse, P., 2008, ‘Mammary tissue damage durin bovine mastitis: Causes and control’, Journal of Animal Scienc 86, 57–65. 10.2527/jas.2007-0302 (http://dx.doi.org/10.2527/jas.2007-0302)

